# General Practitioners’ adherence to prescribing guidelines for statins in the United Kingdoms

**DOI:** 10.1101/625236

**Authors:** Federico Ricciardi, Irwin Nazareth, Irene Petersen

## Abstract

**Objective:** In this retrospective cohort study we aimed to assess, in 202,247 people who started a statin therapy between 2007-2014, the factors that led to the initiation of the drug. To do this we explored CVD risk factors singularly and in combinations as recorded in electronic health records in the year before they receive their first prescription and we compared the risk scores with that suggested by the NICE guideline at that time.

**Methods:** We summarised demographic characteristics and proportions of people with a risk score below the threshold. Regression-based analyses are performed to evaluate the association between the missingness of the risk score and relevant risk score components.

**Results:** 45,364 individuals (22.4%) were prescribed statins without a record of a risk score being available in the year prior to the prescription date. When the risk score was available, 68,174 out of 156,883 patients were prescribed statins even with a score below the 20% threshold. Smoking status was the most frequently recorded variable (74.9% of the instances), followed by systolic blood pressure (71.6%) and total cholesterol (70%), while HDL cholesterol was the least recorded (34.1%). Cholesterol levels are positively associated with the missingness of the risk score, while systolic blood pressure shows a negative association.

**Conclusions:** GPs often start statins on people with no risk score recorded in their clinical records or in those with risk scores below the recommended threshold. Higher cholesterol values may result in a GP starting statin therapy without recording the other relevant components required to calculate a risk score.

**STRENGTHS AND LIMITATIONS OF THIS STUDY:**

- Our cohort contains a large number of individuals: the study provide a representative picture of initiation of statins in UK primary care.
- We specifically focus on which variables and factors GPs record in electronic databases in the year prior statin treatment initiation: this is the first study to directly tackle the issue of statins prescribing in the absence of all the information required by the NICE guideline
- We are not able to verify if GPs actually used the records of the individual health indicators, when these were available, to calculate the risk score.

## 1. INTRODUCTION

Statins are cholesterol-lowering drugs proven to be effective for the primary prevention of cardiovascular disease (CVD)([1]–[6]). The guidance from 2006 published by the UK National Institute for Health and Care Excellence (NICE) recommends statin therapy be initiated in adults aged below 75 years, whose 10-year risk of a cardiovascular diseases (CVD) event is greater than 20%[7], as assessed by “an appropriate risk calculator”[8]. Additionally, the recommendation is that statin therapy be offered to those over 75 years regardless of their 10-year CVD risk score, especially those at high risk of CVD (e.g. smokers or hypertensives). In 2014, NICE updated these guidelines suggesting that those under 75 years of age with a lower risk of 10% CVD risk should be prescribed statins[9]. The risk calculators recommended were the Framingham Risk Score([10]–[13]), and the QRISK2 score[14].

Recent studies have shown that GPs do not adhere to NICE guidelines, such that statins and lipid lowering drugs are prescribed to ineligible patients ([15]–[19]). Non-adherence to the national guidelines is common amongst general practitioner. Recently qualified physicians, however, are more likely to follow guidelines, but this varies by general practice ([20]–[22]). The lack of compliance with the guidelines is twofold: first GPs may prescribe statins even if the risk scores are lower than the recommended threshold and second statins are prescribed without a risk score or its key components being recorded in the clinical records prior to initiation of treatment.

We aim to assess the factors that determine the initiation of statin therapy in people from 2007-2014. In those prescribed statins we:

a. explored which cardiovascular risk factors singly and in combinations are recorded in patient electronic health records in the year before they receive their first prescription;
b. estimated and compared the 10-year CVD risk scores with that suggested by the prevailing NICE guidelines at that time;
c. explored the characteristics of those with and without an available risk score.

## 2. METHODS

### 2.1 Data Source

We used data from The Health Improvement Network (THIN) database (www.thin-uk.com), a large primary care database that provides anonymised longitudinal general practice data on patients’ clinical and prescribing records from more than 500 general practices across the United Kingdom. Over 98% of the UK population are registered with a GP[23] and the database is broadly representative of the United Kingdom population[24]. The database also includes information on the Townsend score, a measure of social deprivation, categorised into five quintiles from one (least deprived) to five (most deprived). Diagnoses and symptoms are recorded by practice staff using Read codes, which is a hierarchical coding system including more than 100,000 codes([25], [26]). The Read code system can be mapped to ICD-10, but in addition the Read codes includes a number of symptoms and administrative codes[26].

### 2.2. Risk Scores

Some individuals had a risk score recorded in THIN and we included this information irrespective of whether these were derived from Framingham or the QRISK2 algorithms. For those for whom this was not recorded, we used any information of the components of this score entered in the previous year to estimate a Framingham score[27]. We use the term “*combined* risk score” for those recorded by the GP or those estimated by us, with the former having precedence if both values were available. In individuals who had with multiple such entries, we considered the one closest to the date of statin initiation.

### 2.3. Study participants

We used data from the time when practices were using their computer system to an acceptable level (ACU date), those in which mortality is satisfactorily recorded (AMR date)([28], [29]) and more than 80% of the patients had a record of social deprivation. We excluded individuals who had a CVD event at any time before statin initiation, or individuals with less than 12 months of data available before treatment initiation date as some of these may not be true incidence cases of statin therapy. We also excluded those without a valid record of Townsend score. Consistently with the NICE guidelines, we excluded people aged 75 or more at the time of statin initiation. Finally, we replaced potential outliers on systolic blood pressure, total or HDL cholesterol, with missing values. Outliers were defined according to a recently proposed two-stage method[30]. In total 781 (0.39%) of outlier values were recorded for any of the variables of interest for this study.

### 2.4. Statistical Analysis

We conducted a descriptive analysis of socio-demographics characteristics and data available within 12 months of first statin prescription for the study population. We identified all those with recorded Framingham Risk Score in the year before first statins prescription, those for whom the risk score has not been recorded but the relevant components were available and those where none of the above were available. We then explored the trends in the recording of each risk factor, by itself and in combination with others. We also obtained descriptive statistics of relevant risk score components (i.e., SBP, total and HDL cholesterol and current smoking status) based on whether the risk score was recorded or not.

We then performed regression-based analyses to evaluate the association between the missing risk scores and the abovementioned risk score components. We fit separate regression models using each one of the four partially observed risk score components in turn as response variable and missingness of the risk score as (i.e., the binary variable taking value 1 when the risk score is missing and 0 when it is available) as independent explanatory variable. When fitting these models, we adjust for the fully observed risk score components (i.e., age at statin initiation, diabetic status, anti-hypertensive treatment and gender) as well as calendar year of statin initiation, to control for known source of confounding. Data extraction and manipulation were done in STATA 13, while data analysis was undertaken in R (version 3.3.1).

## 3. RESULTS

Demographic details of the cohort are given in Table 1. The clinicians started statins on 202,247 individuals, 112,857 (55.8%) men (mean age 58.8) and 89,390 (44.2%) (mean age 60.5) for primary prevention of CVD from 2007 and 2014.

**Table 1.** Demographics of the cohort.

|  | Female | Male | Total |
| --- | --- | --- | --- |
| **Number of subjects** | 89,390 | 112,857 | 202,247 |
| **Age, mean (sd)** | 60.5 (9.0) | 58.8 (9.3) | 59.6 (9.2) |
| **Townsend Score quintile, n (%)** |  |  |  |
| 1 (least deprived) | 22,773 (25.5) | 31,465 (27.9) | 54,238 (26.8) |
| 2 | 20,026 (22.4) | 26,065 (23.1) | 46,091 (22.8) |
| 3 | 18,844 (21.1) | 23,103 (20.5) | 41,974 (20.7) |
| 4 | 16,267 (18.2) | 18,979 (16.8) | 35,246 (17.4) |
| 5 (most deprived) | 11,480 (12.8) | 13,245 (11.7) | 24,725 (12.2) |
| **Statins initiation year, n (%)** |  |  |  |
| 2007 | 13,341 (15.5) | 17,405 (15.4) | 31,246 (15.5) |
| 2008 | 14,389 (16.1) | 18,003 (16.0) | 32,392 (16.0) |
| 2009 | 13,485 (15.1) | 18,274 (16.2) | 31,759 (15.7) |
| 2010 | 10,976 (12.3) | 14,239 (12.6) | 25,215 (12.5) |
| 2011 | 9,659 (10.8) | 12,271 (10.9) | 21,930 (10.8) |
| 2012 | 10,024 (11.2) | 12,466 (11.1) | 22,490 (11.1) |
| 2013 | 9,202 (10.3) | 10,899 (9.7) | 20,101 (9.9) |
| 2014 | 7,814 (8.7) | 9,300 (8.2) | 17,114 (8.5) |

The mean combined risk score was 29.1% (SD 14.6) for men and 18.4% (10.7) for women (Figure **1**). Of those with a risk score, 43,793 (63.6%) women and 24,381 (27.7%) men had a risk score that was lower than the NICE guidelines threshold at that time (i.e. 20%). The risk score distribution was stable over the study period (Table 2).

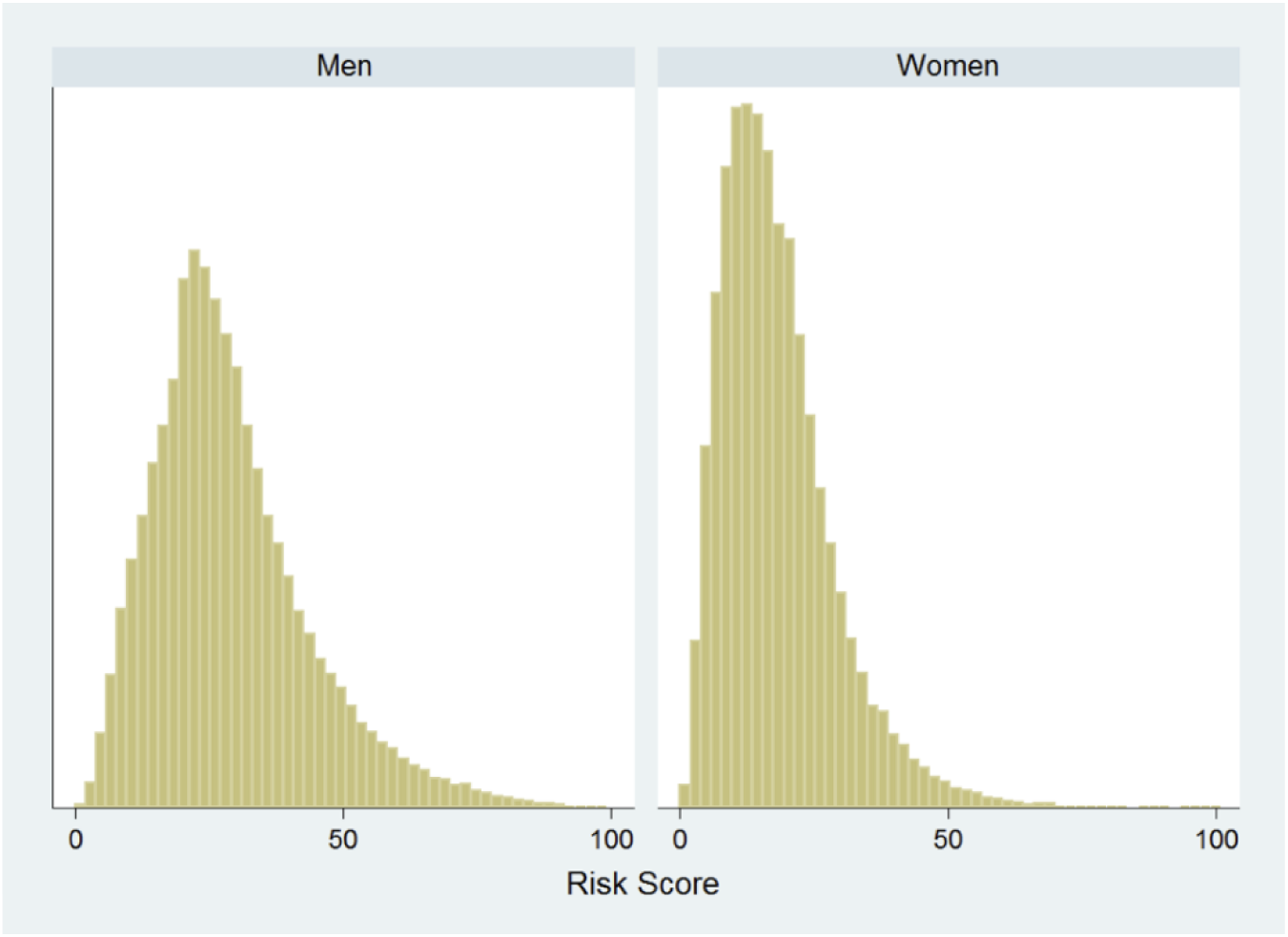

**Figure 1.**
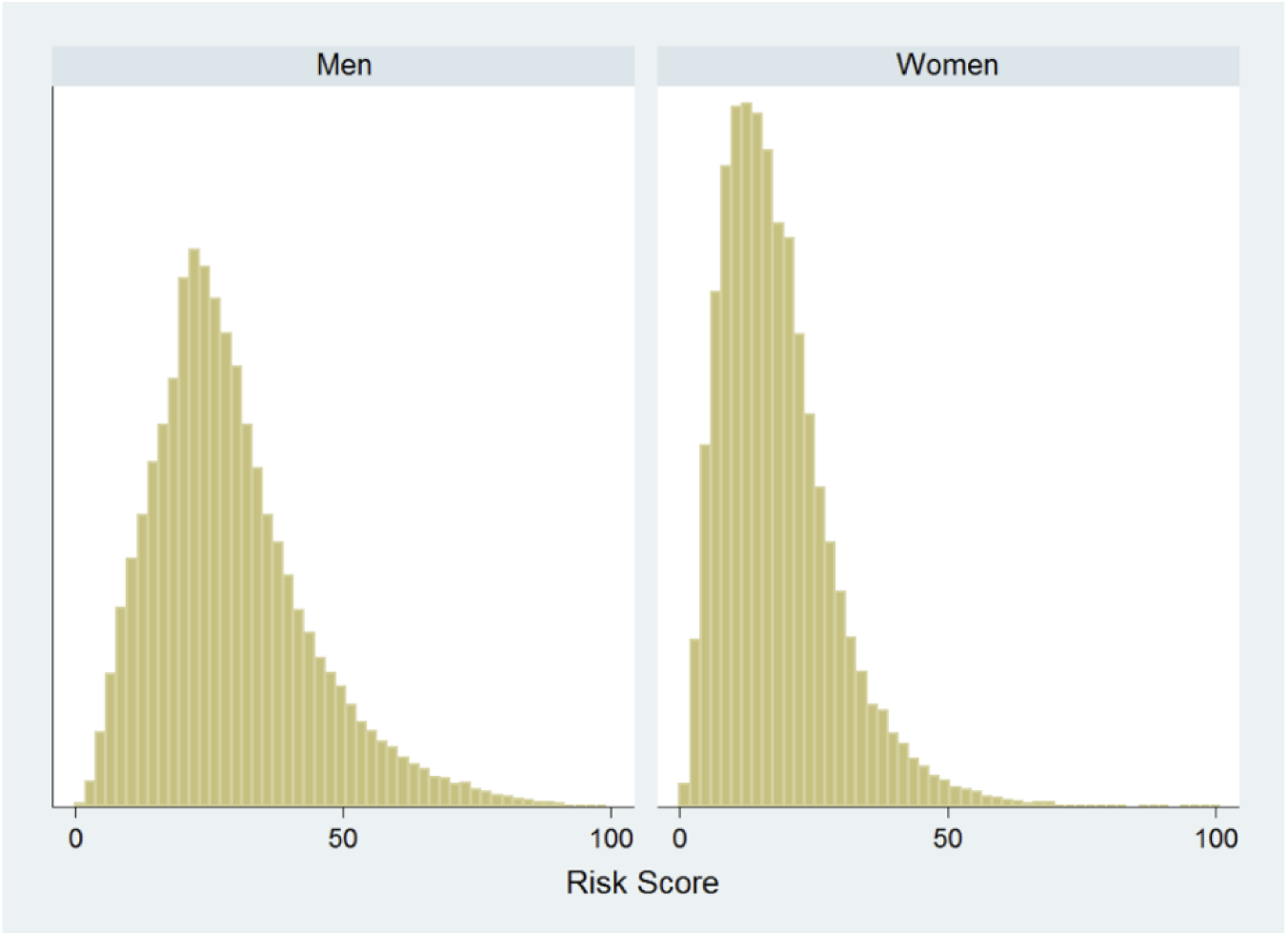
Histograms for the Risk Score, by gender.

**Table 2.** Mean and standard deviation of the Risk Score, by gender and year of statin initiation.

|  | Female | Male | Total |
| --- | --- | --- | --- |
| **Risk Score (%), mean (SD)** |  |  |  |
| 2007 | 18.8 (11.1) | 30.4 (15.4) | 25.3 (14.8) |
| 2008 | 18.5 (10.7) | 29.7 (14.7) | 24.7 (14.2) |
| 2009 | 18.1 (10.5) | 29.3 (14.6) | 24.6 (14.1) |
| 2010 | 18.4 (10.5) | 28.8 (14.3) | 24.3 (13.9) |
| 2011 | 18.2 (10.5) | 29.0 (14.3) | 24.3 (13.9) |
| 2012 | 18.0 (10.6) | 28.3 (14.3) | 23.8 (13.8) |
| 2013 | 18.7 (10.9) | 28.3 (14.3) | 23.8 (13.8) |
| 2014 | 17.9 (10.6) | 27.7 (14.6) | 23.2 (13.8) |
| **Total** | 18.4 (10.7) | 29.1 (14.6) | 24.4 (14.1) |

The likelihood of a missing risk score varied from 2007 to 2014 when stratified by age, gender, Townsend score quintiles, diabetic status and anti-hypertensive treatment. The probability of a missing risk score decreased rapidly after 2008 and was stable thereafter (Figure **2**). Of the 45,364 individuals where a risk score was unavailable, we found that 15,459 (34.1%) of them had a valid record for HDL cholesterol (and in 9,195 (59.5%) cases this was greater than 2); 31,744 (70%) had a valid total cholesterol, of which 2,440 (7.7%) recorded a value lower than 5.

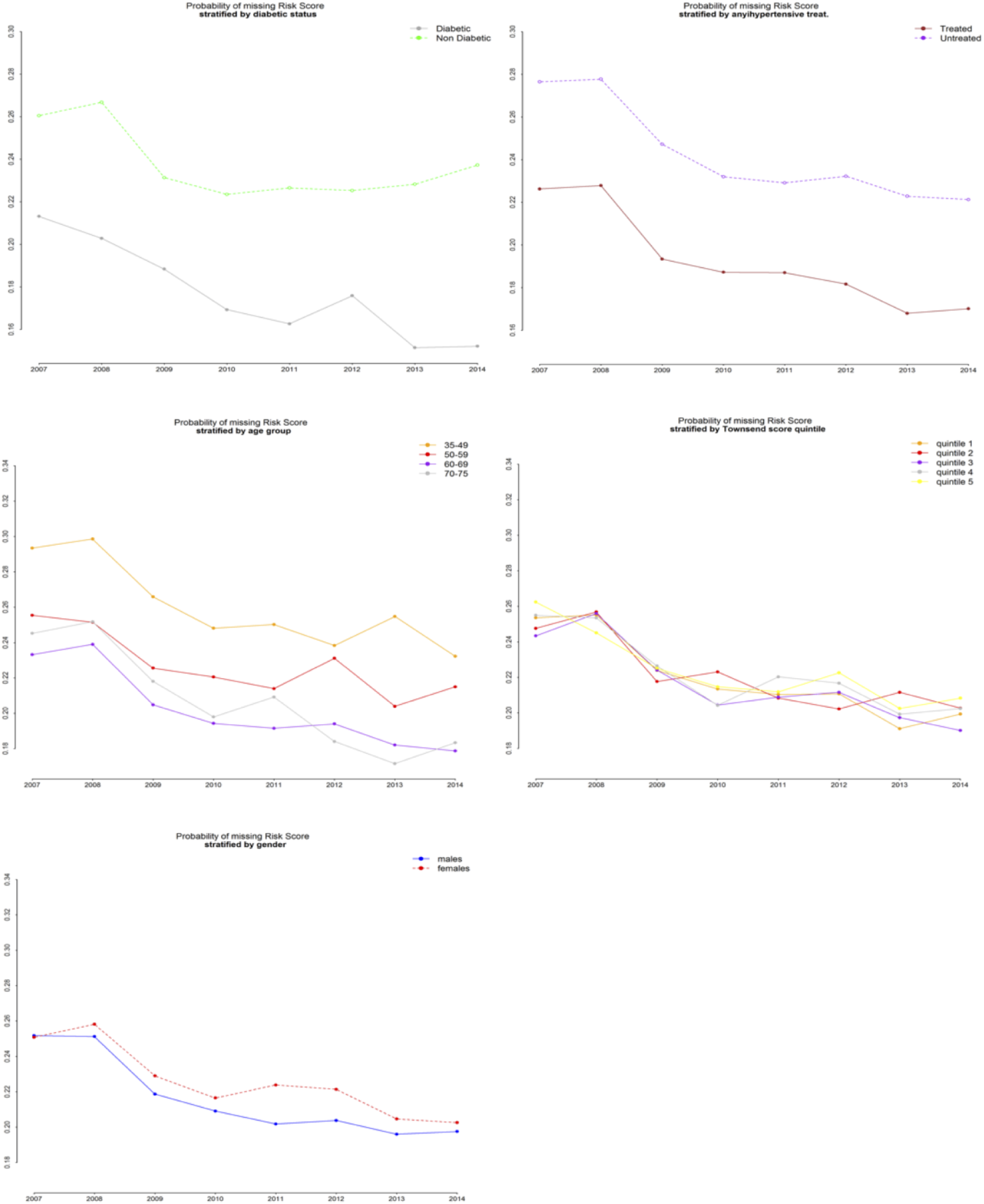

**Figure 2.**
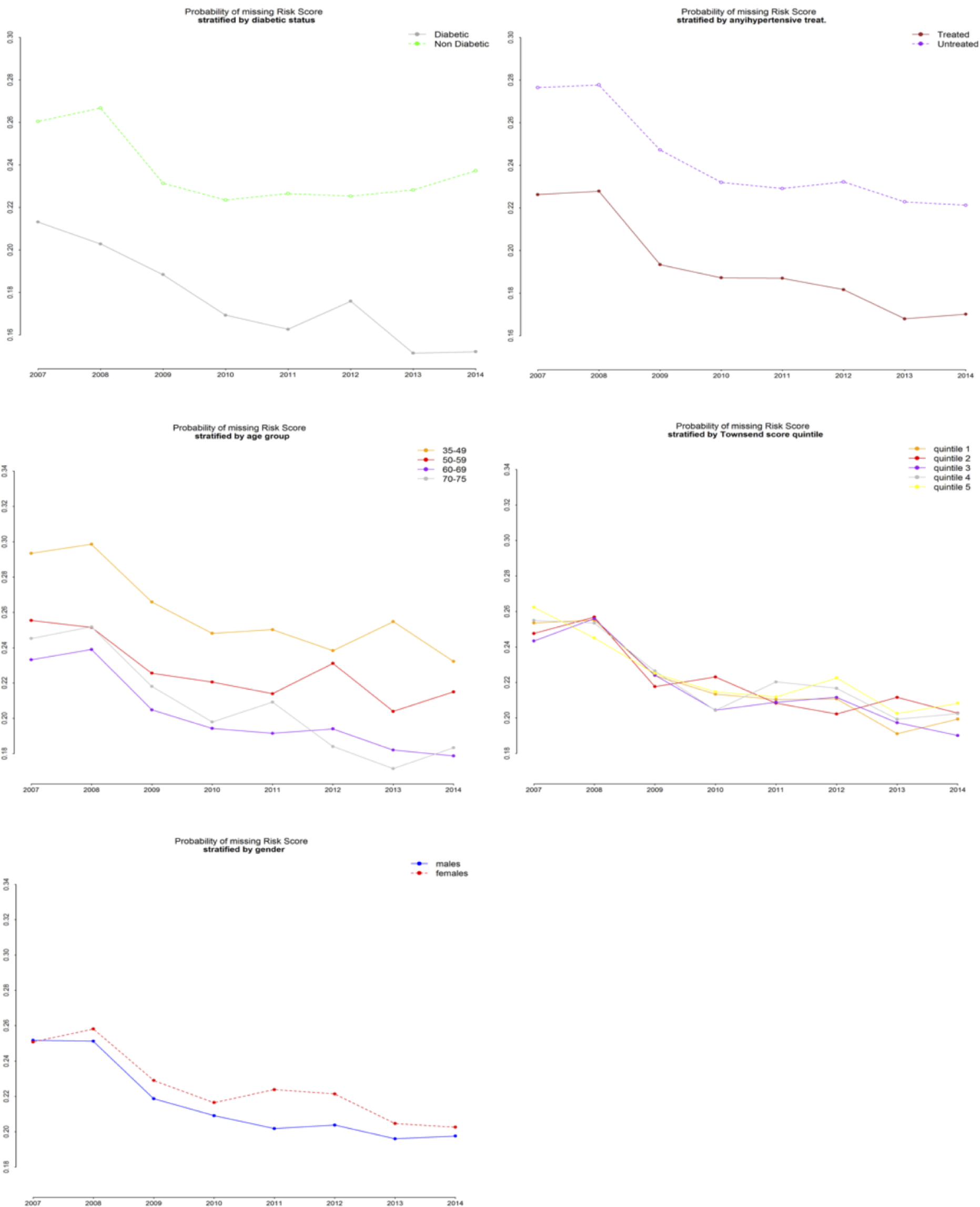
Probability of missing risk score by Year and other characteristics.

Smoking status was the most frequently recorded variable (74.9% of the cases), while HDL cholesterol was the least recorded (34.1%). There were 21,929 (48.3%) individuals without a risk score, who had a record of systolic blood pressure and total cholesterol (Table 3).

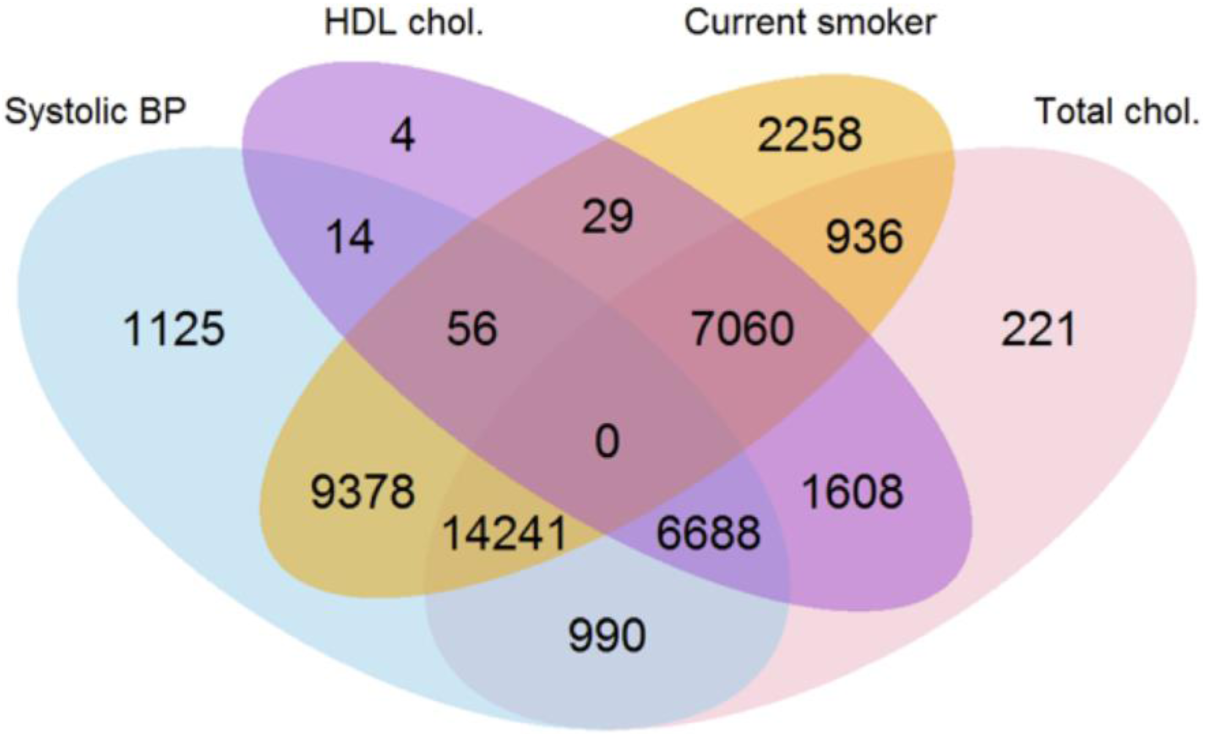

**Table 3.** Observed components for patients with missing risk score. Frequencies for recorded values for each variable are displayed in the main diagonal, while those for jointly recorded values for each pair of variables are given in the cells below.

|  |  |  |  |  |
| --- | --- | --- | --- | --- |
| **SBP** | 32,492<br>(71.6%) |  |  |  |
| **Smoking Status** | 23,675<br>(52.2%) | 33,958<br>(74.9%) |  |  |
| **Tot. cholesterol** | 21,919<br>(48.3%) | 22,237<br>(49.0%) | 31,774<br>(70.0%) |  |
| **HDL cholesterol** | 6,758<br>(14.9%) | 7,145 (15.8%) | 15,356<br>(33.9%) | 15,459 (34.1%) |
|  | **SBP** | **Smoking Status** | **Tot. cholesterol** | **HDL cholesterol** |

**Table 4.** Descriptive statistics (mean, SD and proportion according to the nature of the variable of interest) of the components showing missing values in the two groups defined by the availability of the risk score.

| Variable | Risk score | Mean | SD |
| --- | --- | --- | --- |
| **Total chol.** | available | 6.34 | 1.11 |
|  | missing | 6.47 | 1.16 |
| **HDL chol.** | available | 1.35 | 0.40 |
|  | missing | 1.37 | 0.42 |
| **SBP** | available | 139.41 | 17.05 |
|  | missing | 139.50 | 18.09 |
| Variable | Risk score | % |  |
| **Current smoker** | available | 27.5 | - |
|  | missing | 28.7 | - |

Figure **3** provides details of the various combinations of the risk score components recorded. This offers an overall perspective on recording within the database: the numbers in the overlapping areas represent the observed frequencies of any possible combination of components recorded. For example, of those who did not have a risk score recorded or all the components within a year prior to statin initiation, there were 14,241 (31.4%) who had systolic blood pressure, total cholesterol and smoking status recorded, while 28,045 (61.8%) had a combination of three out of four variables recorded. On the other hand, 2,258 (5%) patients only had smoking status recorded and 1,125 (2.5%) only SBP. Finally, a small amount of 1,647 patients (3.6%) had none of the four variables recorded prior to statins initiation. Summary statistics of the four partially observed components in the two groups defined by the availability of the risk score are given in The levels of both Total and HDL cholesterol was significantly higher among individuals with a missing risk score. However, the value of systolic blood pressure was negatively associated with a missing risk score. Finally, the association between smoking status and a missing risk score was not significant (**Table 5**).

**Figure 3.**
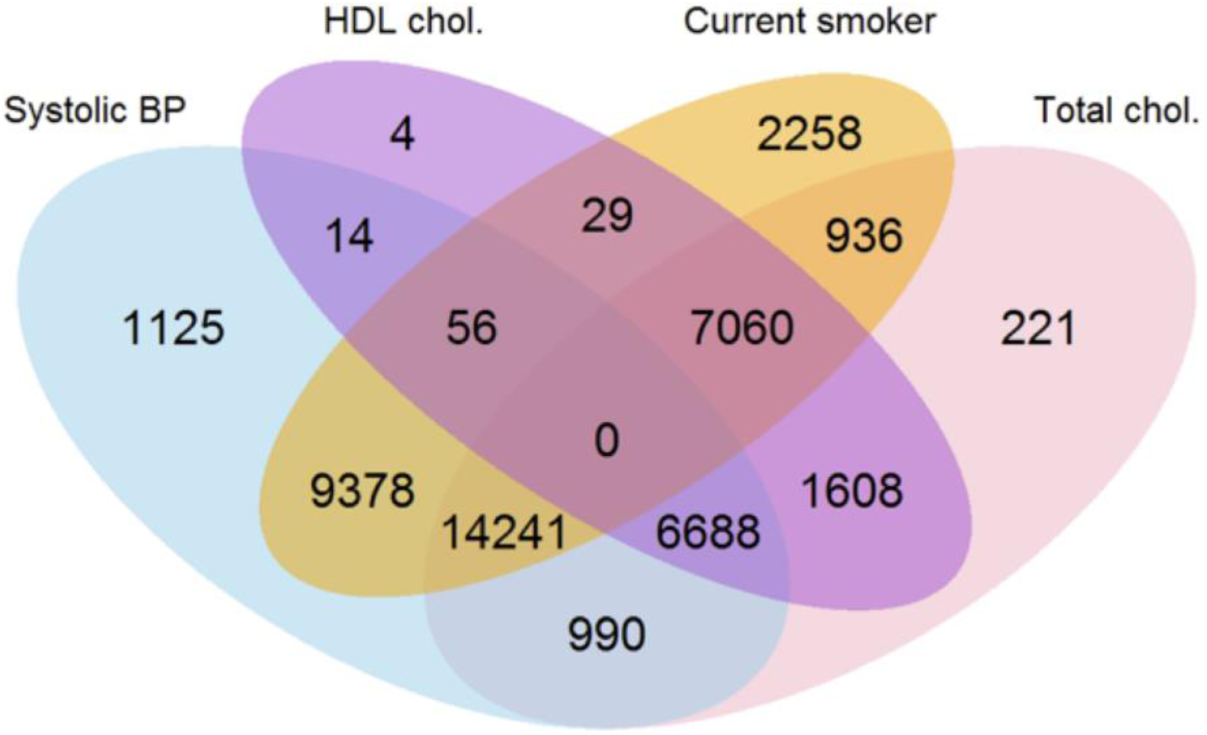
Joint and marginal observed frequencies of the components for patients with missing risk score.

**Table 5.** Results from the regression-based analysis: adjusted means ads standard errors of the components showing missing values in the two groups defined by the availability of the risk score (columns 2-4) and coefficients of binary variable “missing risk score” for listed response variables (columns 5-6), adjusting for fully observable covariates. *\*Odds Ratio is given as coefficient for binary response variable “Current smoker”.*

| **Variable** | **Risk score** | **Adj. mean** | **SE** | **Coefficient** | **95% CI** |
| --- | --- | --- | --- | --- | --- |
| **Total chol.** | available | 6.35 | .003 | 0.054 | (0.041; 0.066) |
|  | missing | 6.41 | .006 |  |  |
| **HDL chol.** | available | 1.35 | .0009 | 0.024 | (0.017; 0.030) |
|  | missing | 1.37 | .003 |  |  |
| **SBP** | available | 139.47 | .042 | -0.295 | (-0.494; -0.096) |
|  | missing | 139.18 | .092 |  |  |
| **Variable** | **Risk score** | **%** |  | **Coefficient\*** | **95% CI** |
| **Current smoker** | available | 27.14 | - | 0.998 | (0.972; 1.025) |
|  | missing | 27.10 | - |  |  |

## 4. DISCUSSION

### 4.1 Key findings

We found that more men (56%) than women (44%) were initiated on statins. On average men were started on the drug at a younger age than women. Less than 20% of the sample had a risk score recorded. About a quarter of people started on statins had at least one of the components of the risk score not recorded in the previous year and 43.5% of those with an available combined risk score were prescribed statins even if their risk score was lower than the threshold set by the guidelines at that time (i.e., 20%). This was more evident in women than in men (63.6% of women versus 27.7% in men). The combined risk score was missing largely due to the absence of *one variable of* the risk score components. In particular, total cholesterol levels are recorded even in absence of the risk score - 70% of those without risk score had a total cholesterol record and 92.3% of these patients had total cholesterol levels above the threshold of 5 mmol/L. We found a significant association between cholesterol levels and a missing risk score and a negative association between systolic blood pressure and a missing risk score.

### 4.2. Strengths and limitations

The major strength of this work is that our cohort contains a large number of individuals, so that our study provided a representative picture of initiation of statins UK primary care. A limitation of our study is that we do not know if GPs used the records of the individual health indicators, when these were available to calculate the risk score. It is likely that many GPs may not have estimated the risk score and recorded it in free text or that a more informal evaluation of the risk score informed the GPs decision-making process.

### 4.3. Comparison with existing literature

There are differences between our study and previous research ([16]–[22]). Whilst we only considered patients started on statins, other studies considered cohorts of both treated and untreated individuals. This was largely because these studies explored the factors triggering the prescribing of these drugs. We aimed to gain an insight into GPs’ prescribing behaviour. With this difference, we can draw some parallels between our findings and those previously described. In a small controlled study Mohammed et al.[16], found that although a 10 year CVD risk was not predictive of statins initiation, a total cholesterol value however, higher greater than 7 mmol/L was predictive. Our results concur with these findings. We found that GPs initiated statin therapy in people with higher total cholesterol values without recording all the relevant components required to estimate a risk score. On the other hand, an elevated blood pressure recording which in general is checked more often than cholesterol may stimulate a more detailed screen of all the components of the risk score. It is also possible that some GPs calculated the risk scores, but did not enter the values into the patient records whilst others may have made their decision on prescribing without estimating CVD risk scores.

Wu et al.[17] found that a considerable proportion of statins prescriptions were given to patients who are not eligible for treatment. Finnikin et al.[18] established that low-risk patients represent a significant proportion of all statin initiations and our results confirm this finding. Christian et al.[19] suggested that older and solo general practitioners rated their clinical judgement to be more effective than guidelines when prescribing statins. Ohlsson and Merlo[21] and Ohlsson et al.[22] suggested that prescribing behaviour within a practice is determined by internal policies and guidelines, availability of equipment, etc., all of which can influence prescribing behaviour explaining the variation between practice prescribing. We examined GPs’ adherence to guidelines based on recorded health indicators required to estimate a CVD risk score but in the absence of any GP information on demographics or practices’ policies we cannot directly compare out results to these papers.

### 4.4. Clinical Implications

Our study relates to the period from 2007-2014, when the prescribing threshold for the 10-year CVD risk was 20%. In 2014, NICE reviewed the guidelines for statins prescribing and they lowered the threshold to 10%. We found that this reduction is in line with the GP practice prior to 2014. GPs started statins on many people with a risk lower than 20%. We would suggest further review of primary care data in the coming years to evaluate if GPs adherence to the new threshold is higher than in the past. GPs often prescribed statins without any record of CVD risk scores in the clinical records even though they recorded information on component CVD risk factors in at least 70% of the people started on statins. This practice needs to change to allow for accurate record keeping and to ensure good continuity of care in the long-term management of people at risk of CVD.

## ACKNOWLEDGEMENTS & FUNDINGS

Approval for use of the THIN data was granted by a medical research scientific review committee on 11th July 2016 (reference 16THIN042). The authors received financial support through grant MR/K014838/1 from the UK Medical Research Council.

## CONTRIBUTORS

All authors were involved in the conception of the study. FR drafted the manuscript. IP and FR contributed to the development of the selection criteria and data extraction criteria. FR performed data extraction and analysis. IN provided clinical expertise. All authors read, provided feedback and approved the final manuscript.

## OPEN ACCESS

This is an Open Access article distributed in accordance with the Creative Commons Attribution Non Commercial (CC BY-NC 4.0) license, which permits others to distribute, remix, adapt, build upon this work non-commercially, and license their derivative works on different terms, provided the original work is properly cited and the use is non-commercial. See: http://creativecommons.org/licenses/by-nc/4.0/

